# Unveiling the PHR-centered regulatory network orchestrating the phosphate starvation signaling in Chinese fir (*Cunninghamia lanceolata*)

**DOI:** 10.1101/2024.06.10.598158

**Authors:** Huiming Xu, Lichuan Deng, Xu Zhou, Yifan Xing, Guolong Li, Yu Chen, Yu Huang, Xiangqing Ma, Zhong-Jian Liu, Ming Li, Liuyin Ma

**Affiliations:** Center for Genomics, Fujian Provincial Key Laboratory of Haixia Applied Plant Systems Biology, Haixia Institute of Science and Technology, College of Forestry, Fujian Agriculture and Forestry University, Fuzhou 350002, Fujian Province, PR China; Fujian Academy of Forestry, Fuzhou 350012, Fujian Province, PR China; Key Laboratory of National Forestry and Grassland Administration for Orchid Conservation and Utilization, Fujian Agriculture and Forestry University, Fuzhou 350002, Fujian Province, PR China

**Author notes:** These authors contributed equally to this work (Huiming Xu, Lichuan Deng).

**Keywords:** PHR, SPX, regulatory network, phosphate starvation signaling, Chinese fir

## Abstract

Phosphorus (P) is an essential mineral element for plant growth and is absorbed and utilized in the form of inorganic phosphate (Pi). However, Pi deficiency largely restricts plant growth in forest ecosystems, while the molecular mechanism of Pi deficiency in woody plants remains unclear. Here, we show that PHOSPHATE STARVATION RESPONSE (PHRs) were central regulators of Pi starvation signaling in Chinese fir, a gymnosperm woody plant. Pi deficiency repressed the shoot growth by decreasing the net photosynthesis rate, reducing the size and number of needle leaves, suppressing the plant height, and reducing the biomass accumulation of shoots in Chinese fir seedlings. Thirteen Chinese fir PHRs (ClPHRs) were characterized, which evolved differently from model and angiosperm woody plants. ClPHRs did not respond to Pi deficiency at the transcriptional level, whereas three ClPHRs responded to Pi deficiency by increasing the nuclear/cytoplasmic protein abundance ratio. Four ClPHRs can restore Pi starvation signaling by activating transcription of *AtPHT1;1* and *AtPHT1;4* in the *atphr1* mutant. Notably, ClPHR7, which is evolutionarily distinct from AtPHR1, was the only ClPHR that could respond to Pi deficiency and restore Pi starvation signals. ClPHR7 could also interact with SPX through protein-protein interaction analysis. Thus, the SPX-PHR regulatory module was also present in gymnosperm woody plants, but the exactly responsible proteins were evolutionarily different from those of model plants. In summary, our results revealed the function of the SPX-PHR regulatory module in Pi starvation signaling and provided genetic information for engineered woody plants with high Pi use efficiency.

## Introduction

Phosphorus (P) is an essential mineral element for sessile plant growth^[1,2]^. P is a functional and structural component of nucleic acids, phospholipids of biomembranes, ATP, and substrate or end-product of enzymatic reactions^[1,2]^. P is critical to plant root growth, shoot growth, reproduction, and fruit quality control by maintaining photosynthesis, respiration, cell growth, energy metabolism, and genetic information transfer^[1]^. However, plants mainly absorb and assimilate P in the form of inorganic orthophosphate (Pi) from soils^[3]^. The availability of Pi in soils is limited due to the slow diffusion rate of Pi in soils and easy chemical fixation by cation ions^[1]^. Thus, Pi deficiency is the second most frequent limiting environmental stress for many natural ecosystems, including forest ecosystems^[1,4]^. Using non-renewable chemical phosphate fertilizers not only increases costs but also leads to environmental pollution due to low Pi use efficiency (only 15-25% of chemical Pi fertilizer is absorbed into plants; others are leaching to the environment)^[2]^. Therefore, understanding the Pi deficiency tolerant mechanism of plants is important.

To cope with Pi deficiency, plants have evolved delicate and complex systems to modulate Pi Starvation Responses (PSRs)^[1,5,6]^. PSRs mainly include an increase in root-to-shoot biomass ratio, a change in root system architecture, an enhancement of Pi uptake, and an increase in Pi use efficiency under Pi stress^[2,6]^. The central regulators of PSRs are PHOSPHATE STARVATION RESPONSE (PHRs), which are MYB transcription factors (TFs) containing both MYB coiled-coil (MYB-CC) domain and MYB-like DNA binding domains^[7]^. PHRs bind to the PHR1-binding sites (P1BS, *GNATATNC*) on the PSR gene promoters to regulate their mRNA expression^[7–9]^. Under Pi deficiency, PHRs enhance the Pi absorption of plants by inducing the transcription of PHOSPHATE TRANSPORT 1 (PHT1s) such as *PHT1;1* and *PHT1;4*, which are Pi influx transporters from soils into the roots^[9–11]^. Moreover, double mutation of PHR1 and PHL1 (PHR1-like 1) largely disrupt PSRs: a >70% of RNA expression level decreases of Pi starvation-induced genes (PSI, ∼ 2000 genes), and a > 50% of RNA expression increases of Pi starvation repressed genes (∼ 1800 genes) in *atphr1atphl1* double mutant of Arabidopsis. Similarly, ∼75% of Pi starvation-associated metabolites are not altered in the *atphr1* mutant of Arabidopsis^[12]^. ATAC-seq analysis in Arabidopsis reveals that mutation of PHR1 and PHL2 greatly reduced the Pi starvation-associated differential accessible genomic regions to nearly 30%^[13]^. The regulatory network analysis also unveils that PHR2 is a central regulator of PSRs-associated arbuscular mycorrhizal symbiosis in rice^[14]^. Overall, PHRs are the central regulators of PSRs.

In Arabidopsis and rice, gene expression analyses show that PHRs do not respond to Pi deficiency at the transcriptional level^[7–10]^. Conversely, PHRs are negatively regulated by SPXs (SPXs: Syg1/Pho81/XPR1)^[1,15,16]^. Under Pi-sufficient conditions, SPXs physically interact with PHRs in the presence of inositol polyphosphates, to prevent PHRs from regulating transcription of the downstream PSR genes^[1,17,18]^. Conversely, SPXs do not interact with PHRs, and the PSRs are activated to enhance the Pi tolerance of plants under Pi deficiency^[1,19]^. Thus, the SPXs-PHRs module is essential in the Pi tolerance of Arabidopsis and rice^[1,3]^. In woody plants, the PtoWRKY40-PtoPHL3-PtoPHT1 module has been reported to regulate the PSRs in poplar^[20]^. The apple PHR1 also enhances Pi tolerance by increasing the transcription of purple acid phosphatase^[21]^. Twenty-one PHRs are identified in tea plants, while their roles in PSRs remain unexplored^[22]^. Overall, the roles of the SPXs-PHRs module in regulating woody plants PSR remain elusive.

Chinese fir (*Cunninghamia lanceolate* (Lamb.) Hook) is a gymnosperm woody plant that belongs to the *Cunninghamia* genus from the *Cupressaceae* family under the Conifer class^[23]^. Chinese fir is native to southern China, Vietnam, and Laos, and is one of the major forest plantation species in southern China and northern Vietnam due to its high economic and ecological values^[23]^. Chinese fir accounts for ∼21.40% of forest plantation areas and ∼30% of wood production from all forest plantations in China^[23]^. However, Pi deficiency is the major limiting environmental stress factor for Chinese fir plantation, primarily due to the high aluminized and acidic soils (pH 4.5-6.4, the active Al^3+^ to fix the Pi into unusable Al-P is 0.8-2.4 mg/g), and the longer Pi recycling cycle from leaves as the Chinese fir leaves do not fall after withering^[23]^. However, how the Chinese fir responds to Pi deficiency via Pi starvation signaling has not been explored.

In this study, we identified that the PSR of Chinese fir required the SPXs-PHRs module. PHRs did not respond to Pi at the transcriptional level, but three PHRs, including ClPHR7, could increase their protein abundance in the nucleus compared to the cytoplasm upon Pi deficiency. ClPHR7 could regulate the PSR by recovering the downstream gene transcription of Pi signaling in the *atphr1* mutant. We further suggest that ClPHR7 could interact with SPXs from a protein-protein interaction network analysis. Notably, ClPHR7 did not evolutionarily belong to the subgroup of AtPHR1. These results suggest that PHRs-SPXs modules were presented in woody plants but do have differences with model plants.

## Results

### Phosphorus deficiency limits the growth of Chinese fir seedlings

To understand the importance of phosphate mineral nutrients in the growth and development of Chinese fir, we analyzed the phenotypes of Chinese fir seedlings upon phosphorus deficiency using hydroponic experiments. In the hydroponic experiments, two-week-old Chinese fir seedlings were divided into two groups: control (0.5 mM KH_2_PO_4_, hereafter referred to as +P) and Pi deficient (0.005 mM KH_2_PO_4_, hereafter referred to as -P), and were grown separately for two months to compare the phenotypic differences between them. Phosphorus deficiency did affect the growth and development of Chinese fir seedlings (Fig. 1). Pi significantly reduced shoots’ fresh and dry weights, while it had little effect on roots (Fig. 1A-E). Therefore, the root/shoot fresh and dry weight ratios were significantly increased in Chinese fir (Fig. 1F-G). Overall, Pi deficiency suppressed the shoot growth of Chinese fir seedlings.

**Fig. 1.**
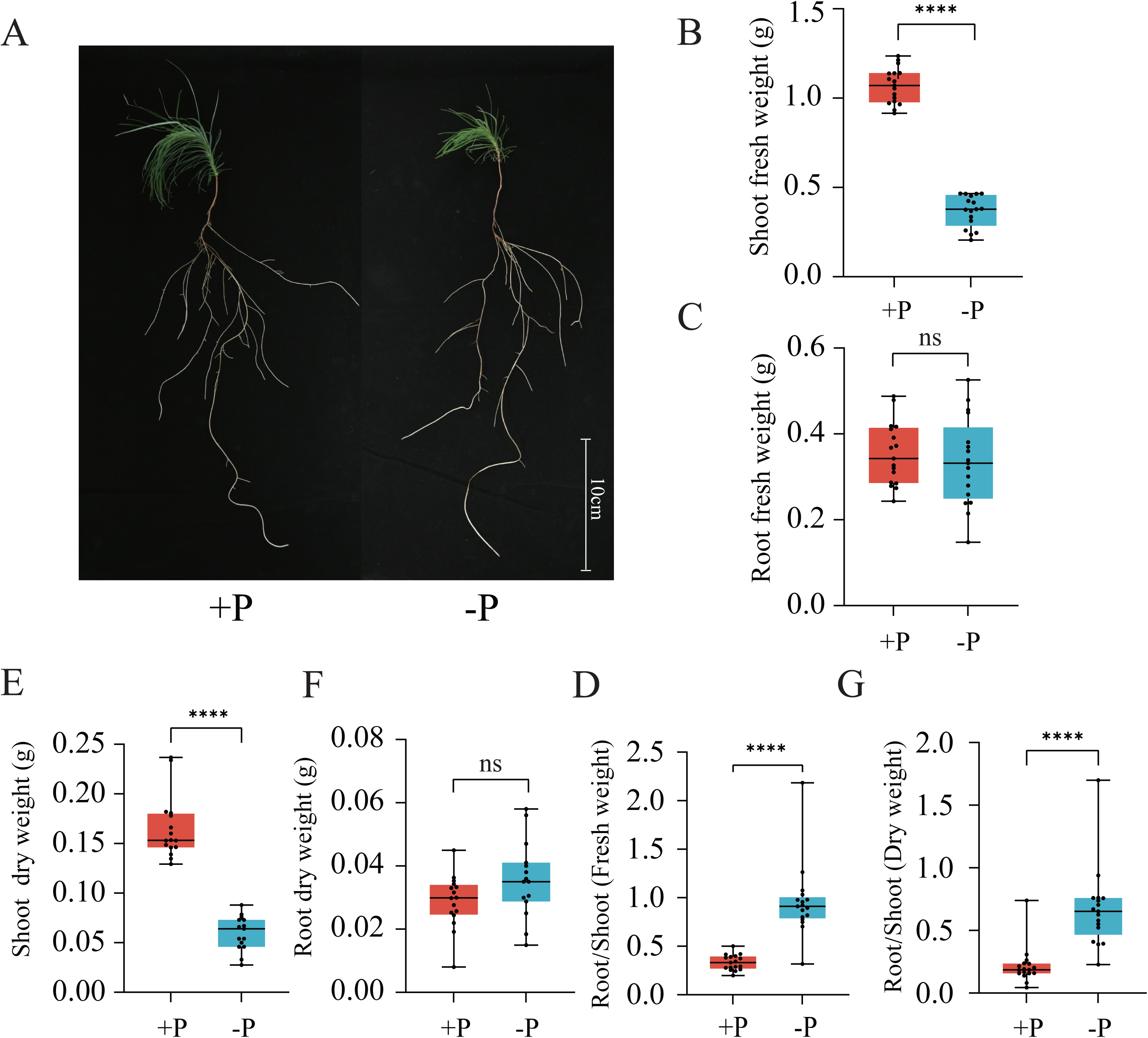
Phosphate deficiency limited the shoot biomass accumulation of Chinese fir seedlings. (A) The growth phenotype of two-week-old Chinese fir seedlings under two-month Pi deficiency treatment (-P, 0.005 mM KH_2_PO_4_) or control (+P, 0.5 mM KH_2_PO_4_). The shoot (B) and root (C) fresh weight of Chinese fir seedlings after two-month growth under -P or +P conditions. The shoot (D) and root (E) dry weight of Chinese fir seedlings under -P or +P conditions for two month. The root-to-shoot ratios of fresh (F) and dry (G) weight upon Pi deficiency in Chinese fir seedlings. “ ns ” represents no significant *P* > 0.05, “ * ” represents *P* ≤ 0.05, “ ** ” represents *P* ≤ 0.01, “ *** ” represents *P* ≤ 0.001, “ **** ” represents *P* ≤ 0.0001.

To further investigate the morphological responses of Chinese fir shoots adapting to Pi deficiency, we separately selected a representative Chinese fir seedling from each control and Pi deficient group to analyze the effects of Pi deficiency on the size and number of Chinese fir needle leaves (Fig. 2A). Pi deficiency also reduced the size and number of Chinese fir needle leaves (Number of needles: +P: 70; -P: 51. Fig. 2A). Notably, the plant height of Chinese fir seedlings was also suppressed upon Pi deficiency (Fig. 2B). Therefore, Pi deficiency reduced the growth and development of Chinese fir needle leaves and stems.

**Fig. 2.**
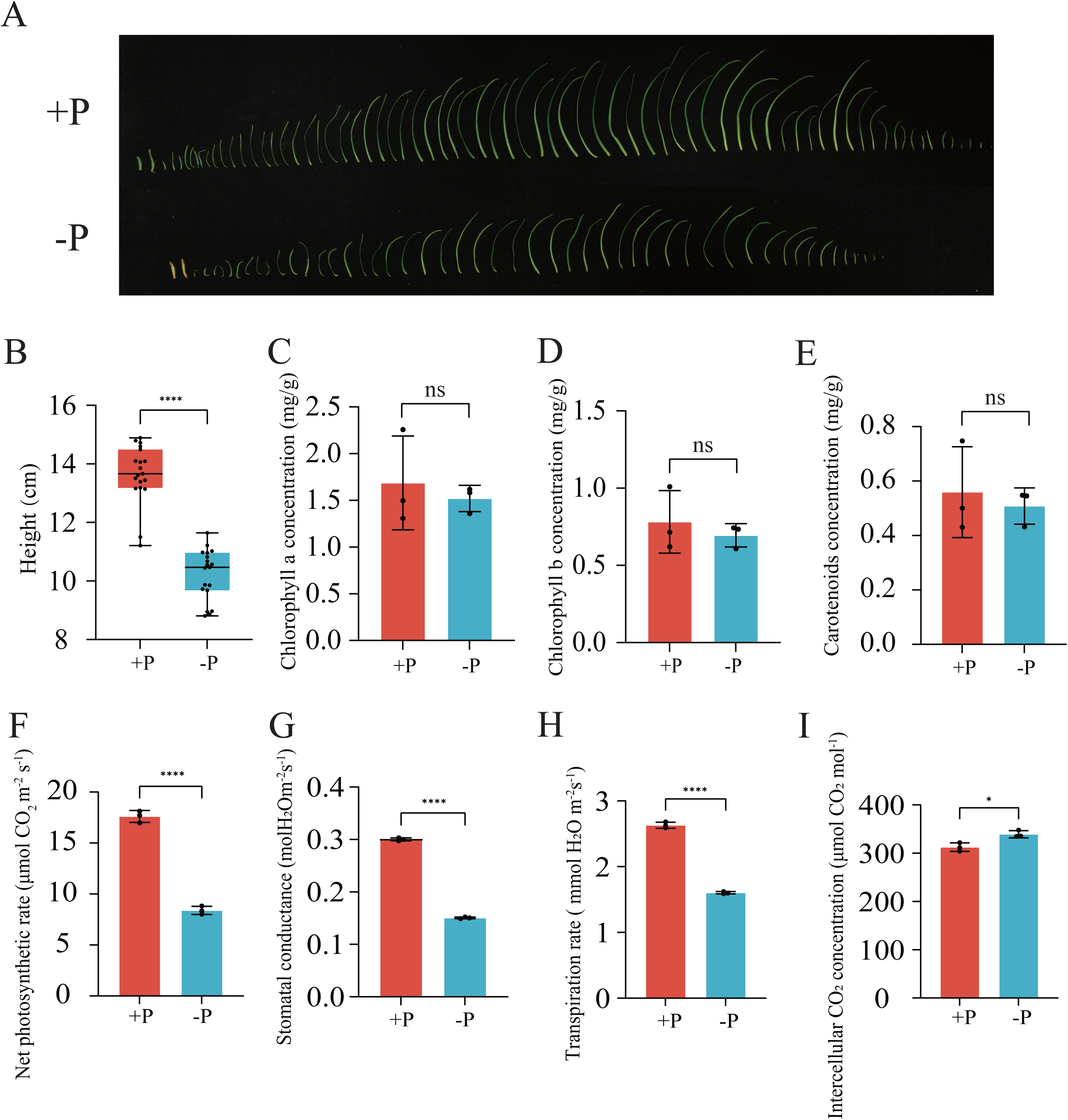
The effect of Phosphate deficiency on the shoot development of Chinese fir seedlings. (A) One-to-one comparison of the needle leaves under Pi sufficient (upper panel) and Pi deficient (lower panel) conditions. The effect of Pi deficiency on plant height (B), the concentration of chlorophyll a (C), the concentration of chlorophyll b (D), the concentration of carotenoid (E), the net photosynthetic rate (F), the stomatal conductance (G), the transpiration rate (H), and the intercellular CO_2_ concentration of Chinese fir seedlings. “ ns ” represents no significant *P* > 0.05, “ * ” represents *P* ≤ 0.05, “ ** ” represents *P* ≤ 0.01, “ *** ” represents *P* ≤ 0.001, “ **** ” represents *P* ≤ 0.0001.

To further explore how the Pi deficiency affects shoot growth and development at the physiological level, we measured the physiological and biochemical indices in Chinese fir needle leaves (Fig. 2C-I). The content of chloroplast pigments (chlorophyll a, chlorophyll b, and carotenoids) of needle leaves did not differ significantly between seedlings in the -P and +P groups (Fig. 2C-E). Therefore, Pi deficiency did not affect the pigment biosynthesis. However, the net photosynthesis rate was significantly suppressed upon Pi deficiency (Fig. 2F). Stomatal conductance and transpiration rate were significantly lower in seedlings in the -P group compared with the +P group, whereas the intercellular CO_2_ concentration in needle leaf cells was significantly higher in Chinese fir needle leaves (Fig. 2G-I). Thus, Pi deficiency reduced the net photosynthetic rate. Overall, these results suggest that Pi deficiency limits the growth and development of Chinese fir seedlings.

### Genome-wide characterization of PHR transcription factors in Chinese fir

Plants adapt to Pi deficiency through PHR-centered systematic Pi signaling^[1]^. To understand the response of PHRs to Pi deficiency, thirteen PHR transcription factors (named ClPHR1∼ClPHR13) were characterized in the Chinese fir genome, all of which contained two domains: Myb_SHAQKYF (PF00249) and Myb_CC_LHEQLE (PF14379) (Fig. S1). The two functional domains of ClPHRs are highly conserved with those of Arabidopsis, poplar, and Chinese pine (Fig. S1). Detailed information on ClPHRs, including the length of CDS, the length of amino acids, the molecular weight of proteins, and the position of chromosomes, is shown in Table 1. The CDS and amino acid sequences of PHRs are listed in Supplementary Tables S1-S3.

**Table 1.** PHR transcription factors in Chinese fir. bp: base pair; MW: molecular weight.

| Name | Gene ID | ORF Length(bp) | Size (aa) | MW (Da) | Location |
| --- | --- | --- | --- | --- | --- |
| CIPHR1 | CI03768 | 1350 | 443 | 48814.24 | 455688859-455717409(-strand) |
| CIPHR2 | CI06722 | 1326 | 435 | 48450.61 | 718374192-718376371(+strand) |
| CIPHR3 | CI14065 | 1320 | 433 | 48475.55 | 1735433-1739082(+strand) |
| CIPHR4 | CI19060 | 1233 | 405 | 46041.69 | 179228692-179231640(-strand) |
| CIPHR5 | CI28538 | 1659 | 545 | 59769.41 | 426573475-426714493(+strand) |
| CIPHR6 | CI32688 | 1125 | 369 | 41498.33 | 610874784-610945634(-strand) |
| CIPHR7 | CI35170 | 1167 | 383 | 42588.87 | 699445289-699488320(- strand) |
| CIPHR8 | CI36481 | 1506 | 494 | 54347.7 | 695731312-695770869(-strand) |
| CIPHR9 | CI12219 | 1098 | 360 | 40854.72 | 2111187-2293968(-strand) |
| CIPHR10 | CI23560 | 1578 | 518 | 56529.66 | 426323558-426368194(+strand) |
| CIPHR11 | CI32526 | 1299 | 426 | 46926.09 | 27608604-27705522(+strand) |
| CIPHR12 | CI22590 | 1266 | 415 | 45289.39 | 13965642-14026212(+strand) |
| CIPHR13 | CI23002 | 651 | 213 | 23911.44 | 186904-190719(-strand) |

The chromosome location of each *ClPHR* was characterized to investigate the genetic differences in the *ClPHR* gene family. A total of 11 chromosomes are characterized in Chinese fir^[23]^. Twelve ClPHRs, except ClPHR13 (located in Contig09589), were unevenly distributed on chromosomes 2, 3, 5, 6, 7, and 11 (Fig. S2). Chromosome 5 is rich in the most abundant members of the *ClPHRs* (*ClPHR2*, *ClPHR5*, *ClPHR9*, and *ClPHR10*), which may indicate that this chromosomal region plays a key role in the duplication and evolution of the *ClPHR* gene family (Fig. S2). Notably, ClPHR3 and ClPHR12, ClPHR5 and ClPHR10, ClPHR8 and ClPHR7 appear in pairs on chromosome 3, chromosome 5, and chromosome 10, respectively (Fig. S2). Overall, ClPHRs were unevenly distributed in the chromosomes of Chinese fir.

### A subgroup of predominantly gymnosperm PHRs was identified in Chinese fir

To understand the potential function of ClPHRs, the phylogenetic analysis was performed among PHRs from Chinese fir, Arabidopsis, Poplar, and Chinese pine. As shown in Fig. 3A, those PHRs can be classified into three subgroups. The first subgroup of PHRs (I) had three ClPHRs (ClPHR8, ClPHR1, and ClPHR12), three Arabidopsis PHRs (including the well-studied AtPHR1), five Poplar PHRs, and three Chinese pine PHRs (PtPHRs). Notably, the second subgroup of PHRs (II) only had PHRs from two gymnosperm species: five ClPHRs (ClPHR5, ClPHR9, ClPHR10, ClPHR11, and ClPHR13) and three PtPHRs (Pt4G66440, Pt3G56740 and Pt4G33370). The last subgroup of PHRs (III) has the most members of PHRs, with five ClPHRs (ClPHR2, ClPHR3, ClPHR4, ClPHR6, and ClPGR7), 10 Arabidopsis PHRs, 16 Poplar PHRs, and seven Chinese pine PHRs. The evolutionary distance analysis of ClPHRs and the other three plants showed that ClPHRs were the most evolutionarily close to the PHRs of Chinese pine, followed by poplar PHRs, and Arabidopsis PHRs were distantly related (Fig. 3A). Overall, these results suggest that gymnosperm PHRs may differ in evolution and function from angiosperm PHRs.

**Fig. 3.**
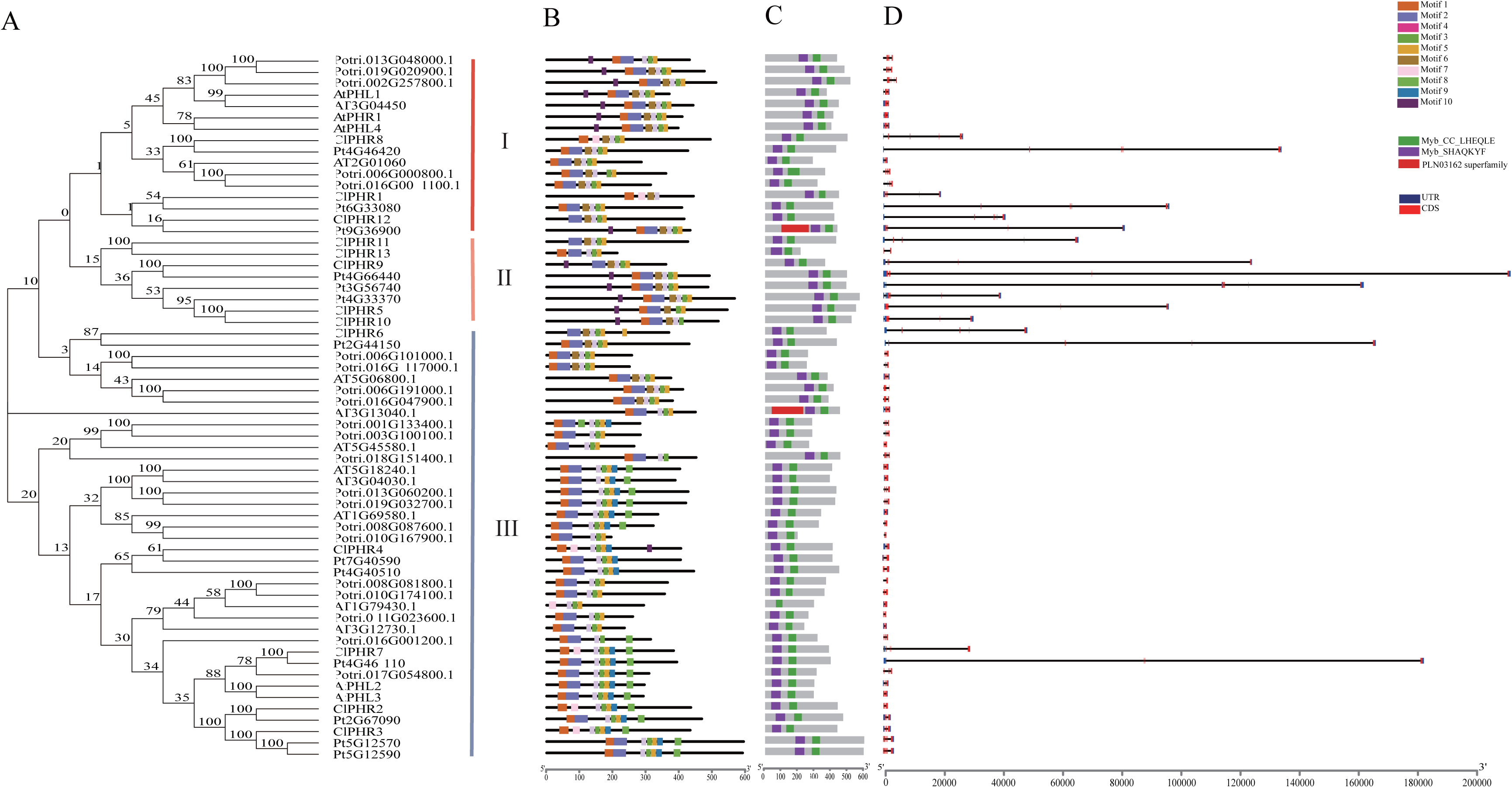
Phylogenetic, functional motifs, and gene structure analyses of Chinese fir PHRs. The phylogenetic (A), conserved motifs (B), conserved structural domains (C), and gene structure (D) analyses of PHRs from Arabidopsis, Chinese fir, Poplar, and Chinese pine.

### ClPHRs were found to be present or absent from specific motifs

To further understand the function of ClPHRs, motif analysis was performed with PHRs from Chinese fir, Arabidopsis, Poplar, and Chinese pine (Fig. 3B). Motif 1, motif 3, motif 5, and motif 6 were present in all PHR proteins, which were annotated as conserved Myb structures (Figs. 3B, S3; Supplementary Table S4). Motif 10 appeared almost exclusively in the first and second subgroups, and Motif 9 only occurred in the third subgroup (Figs. 3B, S3; Supplementary Table S4). Notably, motif 7 (GLTJYHVKSHLQKYRLAKYIPK) was unique to the ClPHRs, and motif 2 (ATPKGVLRVMGVKGLTJYHVKSHLQKYRLAKYLPEESNDKS) is not found in Chinese fir (Figs. 3B, S3; Supplementary Table S4). Therefore, these results suggest that ClPHRs might have different functions than the other three species.

The Conserved domain analysis of PHRs of Chinese fir, Arabidopsis, Poplar, and Chinese pine again showed that all of PHRs possessed two unique domains, namely Myb_SHAQKYF (PF00249) and Myb_CC_LHEQLE (PF14379) (Figs. 3C; S1). However, the gene structure of *ClPHRs* and Chinese pine *PHRs* differed from that of Arabidopsis and Poplar *PHRs* (Fig. 3D). In Chinese fir, nine of *ClPHRs* (all PHRs except *ClPHR2*, *ClPHR3*, *ClPHR4*, and *ClPHR13*), had extremely long introns (average intron length = 7665.2 bp; Fig. 3D; Supplementary Table S5). A similar pattern was also observed in Chinese pine *PHRs* (average intron length = 17777.1 bp; Fig. 3D; Supplementary Table S5). Therefore, compared with the angiosperms Arabidopsis and Populus, the gene structure of gymnosperms *PHRs* was larger and more complex, with extra-long introns.

### ClPHRs did not respond to Pi deficiency at the transcriptional level

To further test the function of ClPHRs, the expression of ClPHRs in different tissues was evaluated. RT-qPCR experiments proved that ClPHRs are expressed in needle leaves, stems, and roots (Fig. S4). However, the expression patterns of different ClPHRs were dynamic in each tissue.

To further investigate their function and regulatory mechanisms, the PlantCARE database was introduced to predict the *cis*-acting elements of *ClPHR*s. A series of *cis*-elements associated with light response, growth and development, hormone signaling, and stress responses were characterized in the promoter regions of *ClPHRs* (Fig. S5). Therefore, ClPHRs are highly regulated under different environmental stress stimuli and are also controlled by different growth and development processes.

Rice OsPHR2 has been reported not to respond to Pi deficiency at the transcriptional level but to respond at the post-transcriptional level^[8]^. In this study, the expression of each *ClPHR* under Pi deficiency at 1 day, 3 days, and 4 weeks was evaluated by RT-qPCR (Fig. S6). Similar to the previous rice results, transcriptional levels of most *ClPHRs* remained largely unchanged upon Pi deficiency (Fig. S6). Therefore, we hypothesize that *ClPHR* may also respond to Pi deficiency at the post-transcriptional level but not at the transcriptional level.

### Four ClPHRs respond to Pi deficiency by changing the nuclear/cytoplasmic protein abundance

To test the hypothesis that ClPHRs may function at the post-transcriptional level in response to Pi deficiency, the expression of ClPHRs with C-terminal GFP fusion was evaluated in the Arabidopsis mesophyll protoplast cells in the presence or absence of Pi. Nine ClPHRs were successfully cloned, and six of them (ClPHR2, ClPHR3, ClPHR6, ClPHR7, ClPHR8, and ClPHR11) were successfully transformed into Arabidopsis mesophyll protoplast cells under +P and -P conditions, respectively (Fig. 4). In rice, OsPHR2 can translocate from the cytoplasm to the nucleus under Pi deficiency to activate the transcription of Phosphate Starvation Induced (PSI) genes such as PHTs^[24]^. Therefore, an increase in the nuclear/cytoplasmic protein abundance ratio upon Pi deficiency can be used as a criterion for screening functional ClPHRs. Notably, the nuclear/cytoplasmic GFP signal intensity ratio of the three ClPHRs (ClPHR3, ClPHR6, and ClPHR7) under the Pi deficiency condition was significantly increased compared to the Pi sufficient condition (Fig. 4). Thus, ClPHR3, ClPHR6, and ClPHR7 might respond to Pi deficiency at post-transcriptional levels. Surprisingly, ClPHR2 reduced the nuclear/cytoplasmic GFP signal intensity ratio under the Pi deficiency condition (Fig. 4), suggesting that ClPHR2 may have the opposite function with ClPHR3, ClPHR6, and ClPHR7 under Pi deficiency. Overall, four ClPHRs responded to Pi deficiency, and three of them might play a positive role in activating the transcription of the PSI genes.

**Fig. 4.**
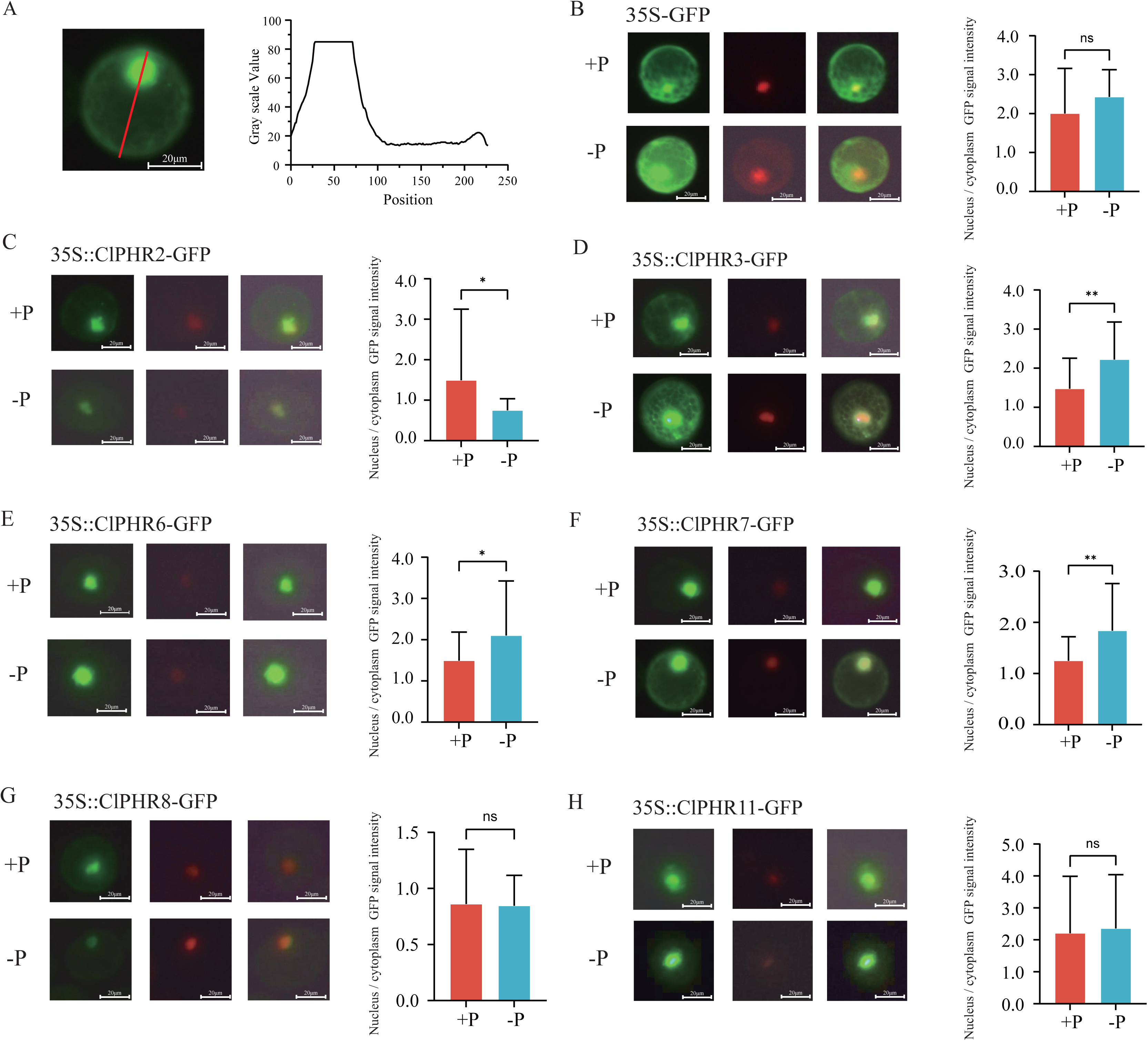
The effect of Phosphate deficiency on the nuclear/cytoplasmic protein abundance ratio of Chinese fir PHRs in Arabidopsis Col-0 protoplasts. (A) The schematic diagram for calculating PHR expression intensity from nuclear/cytoplasmic ratio; The nuclear/cytoplasmic protein abundance ratio of 35S::GFP control (B), 35S::ClPHR2-GFP (C), 35S::ClPHR3-GFP (D), 35S::ClPHR6-GFP (E), 35S::ClPHR7-GFP (F), 35S::ClPHR8-GFP (G), and 35S::ClPHR11-GFP (H) upon Pi deficiency. “ ns ” represents no significant *P* > 0.05, “ * ” represents *P* ≤ 0.05, “ ** ” represents *P* ≤ 0.01, “ *** ” represents *P* ≤ 0.001, “ **** ” represents *P* ≤ 0.0001.

### ClPHRs could recover phosphate starvation signaling

PHRs are the crucial regulators in Phosphate starvation signaling (hereafter refers as Pi signaling) to activate the transcription of downstream PSI genes^[1]^. To evaluate whether ClPHRs were involved in the Pi signaling, the ClPHRs were transformed into the *atphr1* mutant to detect the recovery extent of PSI gene transcription (Fig. 5). Eight of nine cloned ClPHRs except ClPHR8 could successfully recover the expression of *AtPHT1;1* in the mesophyll protoplast cells of *atphr1* mutant (Fig. 5). Notably, the relative expression of *AtPHT1;1* was increased more than 60-fold in the complementary cell line of five ClPHRs (ClPHR2, ClPHR3, ClPHR7, ClPHR10, and ClPHR11; Fig. 5). *AtPHT1;1* and *AtPHT1;4* are two critical high-affinity Pi transporters in Arabidopsis^[8]^. However, only five out of nine ClPHRs (ClPHR2, ClPHR4, ClPHR7, ClPHR10, and ClPHR11) could recover the expression of *AtPHT1;4* in the mesophyll protoplast cells of *atphr1* mutant (Fig. 5). Taken together with three ClPHRs (ClPHR3, ClPHR6, and ClPHR7) that could respond to Pi deficiency (Fig. 4); we speculate that ClPHR7 might play critical roles in Pi signaling as it is the only ClPHR that could respond to Pi deficiency and recover the expression of *AtPHT1;1* and *AtPHT1;4* in *atphr1* mutant.

**Fig. 5.**
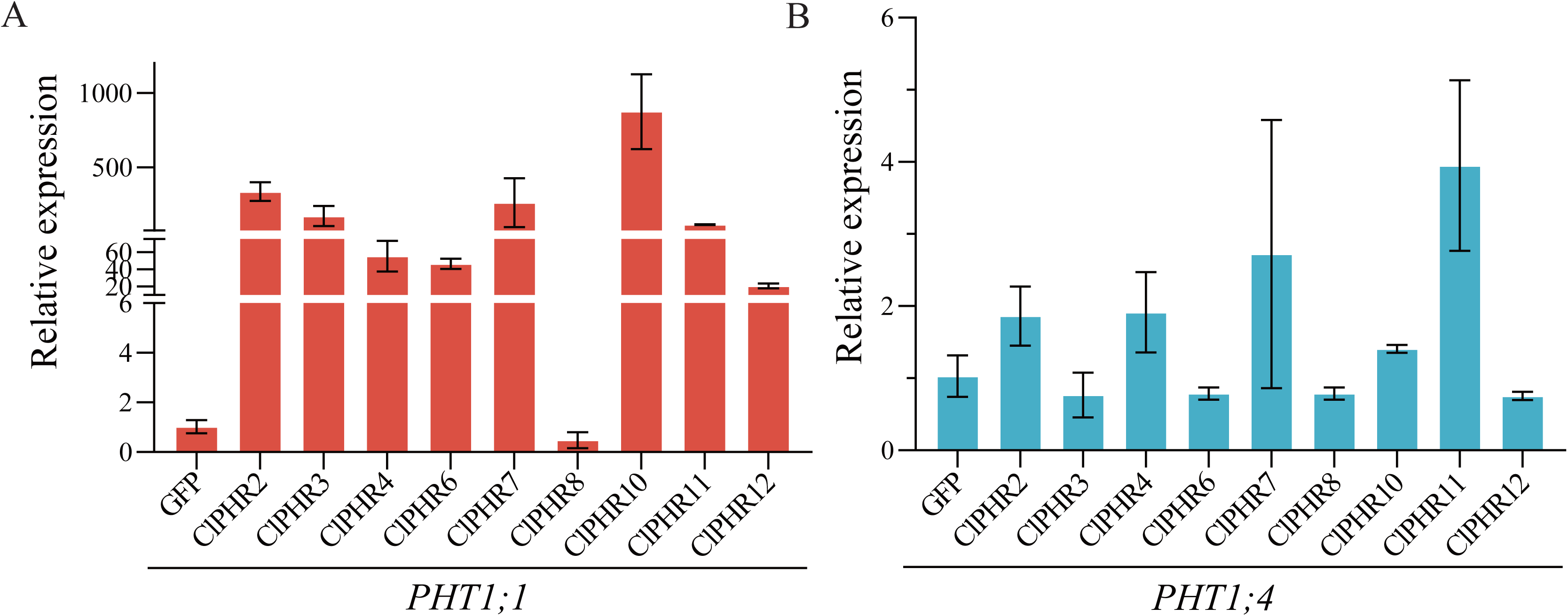
The effect of Chinese fir PHR on the Phosphate starvation signaling in protoplast of Arabidopsis *atphr1* mutant. The relative expression of *AtPHT1;1* (A) and *AtPHT1;4* (B) in the Arabidopsis protoplasts after complementing the Chinese fir PHRs into *atphr1* mutants.

### ClPHRs interacted with SPXs in the Protein-Protein Interaction network analyses

To explore the potential function of ClPHRs, the protein sequences were uploaded to the STRING database to construct the protein-protein interaction network (Fig. 6). The prediction results showed that multiple ClPHRs might be involved in interacting with Pi signaling crucial regulators (Fig. 6). ClPHR7 was at the hub of the protein-protein interaction network. ClPHR7 could interact with AtSPX1 and AtSPX2 (Fig. 8), and these two SPXs play crucial negative roles in regulating the function of PHRs-centered Pi signaling in plants^[1]^. ClPHR7 could also interact with the AtPAP10-2 (Fig. 6), a functional purple acid phosphatase induced under Pi deficiency at the transcriptional and post-transcriptional levels^[25]^. ClPHR7 also interacted with ARD1 and GPA1 (Fig. 6), which are cell growth regulators. Therefore, these results further suggest that ClPHR7 might play a crucial role in regulating Pi starvation signaling. Similar to ClPHR7, ClPHR2, ClPHR10, ClPHR12, and ClPHR13 also interacted with AtSPXs. Overall, these results suggest that ClPHRs might function on Pi signaling regulation via interacting with SPXs.

**Fig. 6.**
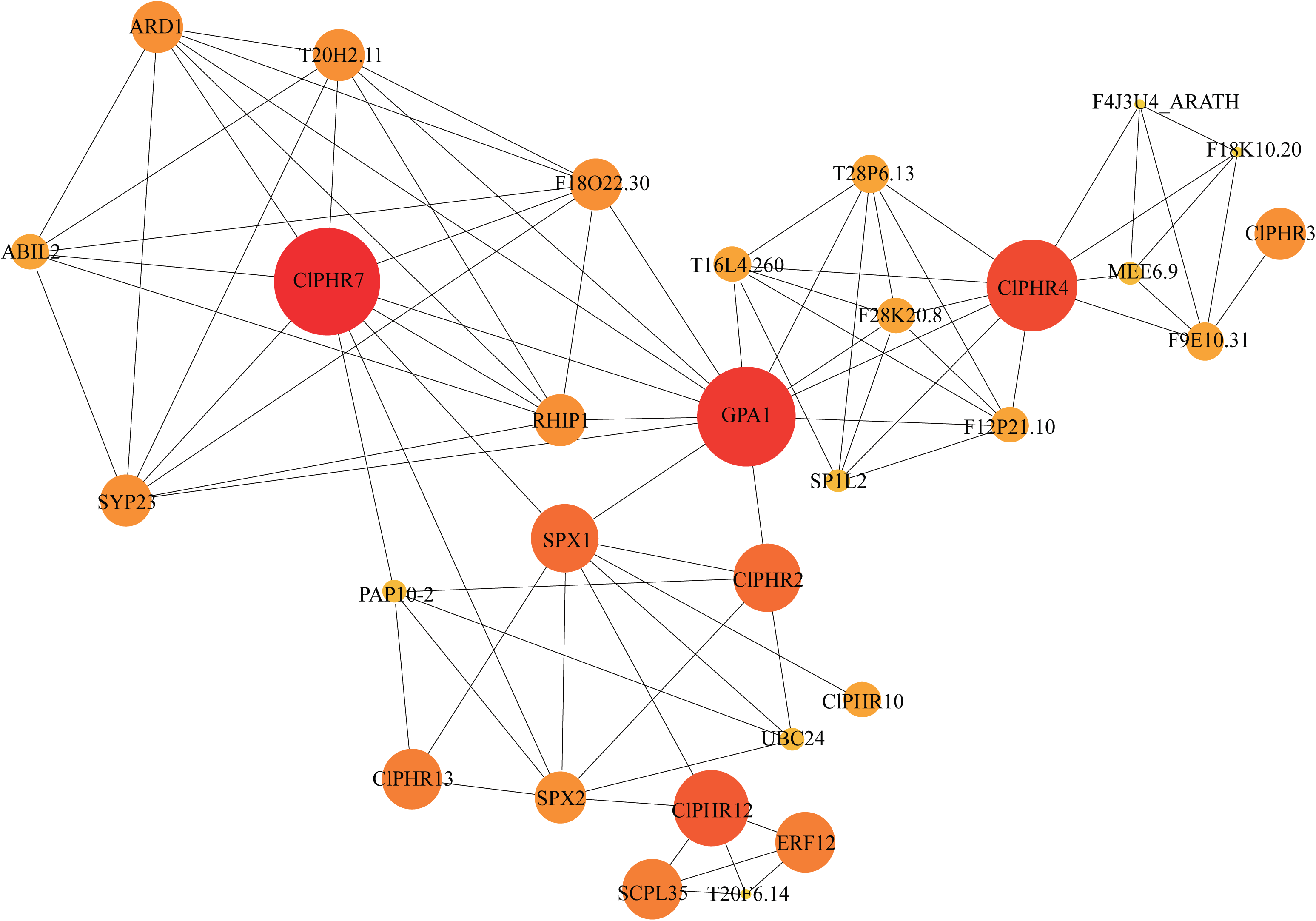
The Protein-protein interaction network of the Chinese fir PHRs. The network diagram consists of nodes and edges, where each node represents a protein. The connections between nodes (edges) represent the interactions among these nodes, and the nodes’ depth of node color and size indicate the nodes’ weight.

## Discussion

Pi deficiency is one of the major environmental stresses in many natural ecosystems, including forest ecosystems^[1,4]^. The underlying molecular mechanisms of Pi starvation signaling have been systematically studied in model and crop plants^[2,5,6]^. However, despite their importance, the molecular mechanisms of Pi deficiency in woody plants have lagged behind compared to the model and crop plants^[1]^. In this study, the PHR-centered molecular mechanism in the gymnosperm woody plant Chinese fir has been characterized.

Similar to the other two woody plants (apple and *Citrus grandis*), Pi deficiency also suppressed shoot growth and reduced the net photosynthesis rate in Chinese fir^[1]^. Pi deficiency reduced the net photosynthesis rate in two potential ways: (1) Pi deficiency impairs the electron transport between photosystem II and photosystem I to reduce the net photosynthesis rate in *Citrus grandis* ^[1]^. (2) The low stomatal conductance reduces the rate of CO2 diffusion and decreases the net photosynthetic rate upon Pi deficiency in wheat^[26]^. Similar to wheat, the stomatal conductance and transpiration rate were significantly lower, whereas the intercellular CO_2_ concentration in needle leaf cells was significantly higher upon Pi deficiency in Chinese fir seedlings (Fig. 2G-I). Taken together, we speculated that the reduced net photosynthetic resulted in lower stomatal conductance in Chinese fir.

Although PHRs have been identified in Arabidopsis, rice, maize, soybean, cotton, and tea plants, functional analyses of PHRs from non-model plants remain largely unexplored^[27–29]^. This study characterized thirteen PHRs in the gymnosperm woody plant Chinese fir. Notably, a subgroup of PHRs only identified in two gymnosperms-Chinese fir and Chinese pine, suggesting that there might be gymnosperm-specific PHRs (Fig. 3). It will be interesting to test the function of these PHRs and understand why they were only dominant in gymnosperm in the future. Similar to the other gymnosperm, the intron length of ClPHRs was far larger than that of angiosperm, primarily due to the extremely larger genome of the gymnosperm (Fig. 3)^[30]^. Overall, our results suggest that gymnosperm PHRs might differ from angiosperm.

The genetic transformation of Conifer remains a challenge^[31]^. To complement the lack of a genetic transformation system in Chinese fir, we have established a complex system to evaluate the function of ClPHRs in Pi starvation signaling. Similar to the results from rice^[8,9,16]^, the RNA expression level of *ClPHRs* did not respond to Pi deficiency (Fig. S6). Therefore, we introduced the ClPHRs in protoplast and characterized ClPHR3, ClPHR6, and ClPHR7 with an increased nuclear/cytoplasmic protein abundance ratio upon Pi deficiency (Fig. 4). To further understand the function of ClPHRs on Pi starvation signaling, we complement the ClPHRs into the protoplast of *atphr1* mutant, and ClPHR2, ClPHR7, and ClPHR10 could recover the RNA expression of *AtPHT1;1* and *AtPHT1;4* (Fig. 5). As only ClPHR7 responded to Pi deficiency and could activate transcription of PHT1;1, therefore, ClPHR7 was a potential key candidate that functions on Pi starvation signaling. In Arabidopsis and rice, PHRs were negatively regulated by SPXs^[3,9]^. Protein-protein interaction analyses proved that ClPHR7 could interact with SPX1, SPX2, and PAP10-2 (Fig. 6). Therefore, we speculated that SPX-ClPHR7 was a hub regulatory module for Pi starvation signaling. Further analyses with Chinese fir protoplast are required to understand the function of the SPX-ClPHR7 regulatory module in Chinese fir.

Overall, unveiling the SPX-PHR module in Chinese fir can reveal the biological functions of PHR transcription factors in woody plants, and also provide a basis for molecular breeding of high Pi use efficiency Chinese fir in the future. In addition, these PHRs can also act as molecular markers to select elite germplasm resources from current Chinese fir germplasm resource stocks.

## Materials and Methods

### Plant materials and treatment

Two types of Arabidopsis plants were used in this study: the *Arabidopsis thaliana* ecotype Columbia wild-type (Col-0) and a T-DNA insertion mutant line of AtPHR1 (AT4G28610), *atphr1* (SALK_067629), for the protoplast transient expression assay ^[32]^. The Chinese fir mixed seeds were collected from Jiangle State Forestry Farm (E117°47□; N26°73□), Jiangle, Fujian Province, China.

10 g Chinese fir seeds were put on vermiculite for germination and growth. The Chinese fir seedlings were grown in a growth chamber with 16 hr light/ 8 hr dark at 23.5 °C. Two-week-old Chinese fir seedlings with similar vigor were selected to grow in phosphorus-sufficient conditions for five days to replenish nutrients. The Chinese fir seedlings were then divided into two groups: + P group (0.5 mM KH_2_PO_4_, hereafter referred to as +P) and Pi deficient group (0.005 mM KH_2_PO_4_, hereafter referred to as -P) for two months. The culture medium was changed every three days. The detailed information on the hydroponic nutrient medium was identical to the previous study (-P group was identical to HNLP; +P group was identical to HNHP) ^[33]^. Similarly, two-week-old Chinese fir seedlings were also grown one day, two days, and four weeks under +P and -P groups for gene expression analyses. Samples from each of the time points have three biological replicates.

### Phenotypic analysis

The Plant Chlorophyll and Carotenoid Content Determination Kit from Shanghai Zhuangcai Biotechnology Co., Ltd. (Cat: ZC-S0886) determined the chlorophyll a, chlorophyll b, and carotenoid content of Chinese fir seedlings with three biological replicates. The net photosynthetic rate, transpiration rate, stomatal conductance, and intercellular CO_2_ concentration were measured by LI-6400 XT Portable Photosynthesis Assay System from Beijing Ecotek Technology Co., Ltd. The height of Chinese fir seedlings was calculated using ImageJ (> 25 individual seedlings).

### Characterization of PHR transcription factors in Chinese fir

To identify the PHR transcription factors in Chinese fir, the Arabidopsis PHR amino acid sequences were downloaded from the TAIR database (https://www.arabidopsis.org/). The Arabidopsis PHRs were then blasted to Chinese fir peptide sequences by Blastp with the criteria (expected value < 1×10^-5^) ^[23,34]^. The candidate Chinese fir PHR transcription factors were then used to predict the functional domain with default parameter by Pfam (version 36.0; http://pfam-legacy.xfam.org/) ^[35]^. Similar to previous studies ^[27,29]^, the Chinese fir PHR transcription factor candidates, which harbored all two functional domains: Myb_SHAQKYF (PF00249) and Myb_CC_LHEQLE (PF14379), were considered as the Chinese fir PHR transcription factors (ClPHRs, Supplementary Table S1-S2). The *Populus trichocarpa* (poplar) and Chinese pine genome, CDS sequences, and amino acids sequences were separately downloaded from PopGenIE ^[36]^ and CNGB sequence Archive: CNP0001649 ^[30]^. The poplar and Chinese pine PHRs were identified using methods identical to Chinese fir PHRs. The CDS, amino acid sequences, and chromosome localization of ClPHRs were extracted from Chinese fir genome annotation ^[23]^. The isoelectric point (PI) and molecular weight (MW) of ClPHRs were calculated by the “Compute pI/Mw” tool available at Expasy (https://web.expasy.org/compute_pi/).

### Bioinformatics analyses

The PHR amino acid sequences of Chinese fir, Arabidopsis, poplar, and Chinese pine were aligned using MEGA7 software, and the phylogenetic tree was constructed using the adjacency method (bootstrap analysis of 2000 repeats)^[37]^. Motif analyses were performed using the MEME suite (version 5.5.5; https://meme-suite.org/meme/tools/meme)^[38]^. The location information of the ClPHRs on the chromosome of Chinese fir was analyzed and visualized by MG2C online tool (version 2.1; http://mg2c.iask.in/mg2c_v2.1/). To predict and generate ClPHR protein interaction networks based on known Arabidopsis homologous proteins, the STRING database (https://string-db.org) was used. The protein-protein interaction map was drawn with Cytoscape (version 3.10.2) ^[39]^.

### Gene expression and gene cloning

The gene expression analyses of ClPHRs under Pi deficiency of one day, three days, and four weeks, as well as the tissue expression pattern of ClPHRs, were performed as described in a previous study by RT-qPCR^[40]^. ACTIN1 and ACTIN2 were used as the internal control, and the expression level of ClPHRs was calculated according to the 2^−ΔΔCt^ method^[40]^. The CDS sequences without stop codon of each ClPHR were separately cloned into a modified pCambia3301 vector^[41]^, to construct the 35S:: ClPHR-GFP construct at SpeI restriction digestion site using a homologous recombination method^[41]^. The Sanger sequence proved plasmids were then used for the transient expression analysis of ClPHRs into Arabidopsis mesophyll protoplast cells^[40,42]^. The primers used in this study were listed in Supplementary Table S6.

### Protoplast transient expression

For functional analyses of ClPHRs in *atphr1* mutants, 5∼20 leaves from 3∼4-week-old Arabidopsis *atphr1* mutants were used for isolating mesophyll protoplast cells according to the previous protocol^[40]^. 100 μg 35S::ClPHRs-GFP, 35S::GFP (as control) constructs were separately transformed into *atphr1* mutants with the PEG method as described in previous study^[40]^. The RNA isolation and gene expression analyses of *AtPHT1;1* and *AtPHT1;4* were identical to the previous study^[40]^.

For the nuclear/cytoplasmic protein abundance ratio analysis under Pi deficiency, Arabidopsis Col-0 was used to isolate the mesophyll protoplast cells. 100 μg 35S::ClPHRs-GFP and 35S::GFP (as control) constructs were separately transformed into the mesophyll protoplast cells with either Pi deficiency WI buffer (-P, 0.005 mM KH_2_PO_4_, 0.5 M D-Mannitol, 4 mM MES, 20 mM KCl) or normal Pi supply WI buffer (+P : 0.5 mM KH_2_PO_4_, 0.5 M D-Mannitol, 4 mM MES, 19.5 mM KCl). The GFP signals were detected by a microscope (20x, Novel, NIB900). The gain value of GFP was set to 2, and the exposure time was set to 350 ms. The nuclear/cytoplasmic ratios of each ClPHR (>30 individual cells) were calculated using ImageJ (https://imagej.net/ij/index.html).

## Author contributions

Liuyin Ma, Ming Li, and Zhong-jian Liu conceived the ideas; Huiming Xu, Lichuan Deng, Xu Zhou, and Guolong Li performed the experiments; Huiming Xu, Lichuan Deng, and Yifan Xing analyzed the data; Yu Chen, Yu Huang, and Zhong-Jian Liu provided the genome sequences and Chinese fir materials. Liuyin Ma, Huiming Xu, Ming Li, and Xiangqing Ma wrote the manuscripts. All authors proved manuscripts.

## Acknowledgements

The authors thank Prof. Wenfei Wang from Fujian Agriculture and Forestry University for kindly providing Arabidopsis *atphr1* mutant. This work was supported by the National Key Research and Development Program of China (2021YFD2201302), the National Natural Science Foundation of China (31971674), the Basic Scientific Research Projects for Provincial Public Welfare Research Institutes from Fujian Provincial Department of Science and Technology (2022R1010003), the Forestry Peak Discipline Construction Project of Fujian Agriculture and Forestry University (72202200205), and the Annual Funding from Center for Genomics at Fujian Agriculture and Forestry University (11899001003).

## Supplementary data

**Supplementary Fig. S1.** Multiple sequence alignment of the two conserved ClPHRs domains. The Myb_SHAQKYF and Myb_CC_LHEQLE domains were marked in the red box. The highlighted color represents the degree of amino acid identity: black represents a sequence identity of 100%; pink represents a sequence identity between 80% and 99%; blue represents a sequence identity between 50% and 79%.

**Supplementary Fig. S2.** Chromosomal localization of the Chinese fir *PHR* genes. The chromosome numbers are shown at the top of each chromosome. The name for ClPHR members marked in red color is shown on both sides of each chromosome, and its position on the corresponding chromosome is shown on the left.

**Supplementary Fig. S3.** Schematic representation the sequence logo of the ClPHR protein motif.

**Supplementary Fig. S4.** The Cis-elements of Chinese fir PHR transcription factor. The four categories of cis-elements are light-response elements, hormone-responsive, stress-responsive, and plant development. Different colors indicate different cis-elements and their ratios present in ClPHR genes.

**Supplementary Fig. S5.** The relative expression of the Chinese fir *PHR* gene in roots, stems, and needle leaves. The relative expression of *ClPHRs* in the root (A), stem (B), and need leaves (C) of Chinese fir four-week-old seedlings. “ ns ”, no significant, represents *P* > 0.05,“ * ” represents *P* ≤ 0.05, “ ** ” represents *P* ≤ 0.01, “ *** ” represents *P* ≤ 0.001, “ **** ” represents *P* ≤ 0.0001.

**Supplementary Fig. S6.** Schematic heatmap represents relative expression of Chinese fir *PHR* genes upon Pi deficiency. The heatmap represents the expression of Chinese fir PHR genes under 1-day, 3-day, and 4-week Pi deficiency or sufficient treatment.

**Supplementary Table S1.** The CDS Sequences of Chinese fir *PHRs*

**Supplementary Table S2.** The amino acid sequences of Chinese fir PHRs

**Supplementary Table S3.** The amino acid sequences of PHRs from Arabidopsis, Chinese pine, and Poplar

**Supplementary Table S4.** The motif sequences of ClPHRs analyzed by MEME

**Supplementary Table S5**. The detailed intron information of PHRs from Arabidopsis, Chinese fir, Chinese pine, and Poplar

**Supplementary Table S6**. List of Primers used in this study

