## Supplementalry Figures for "Unveiling the PHR-centered regulatory network orchestrating the phosphate starvation signaling in Chinese fir (*Cunninghamia lanceolata*)"

Supplementary Fig. S1

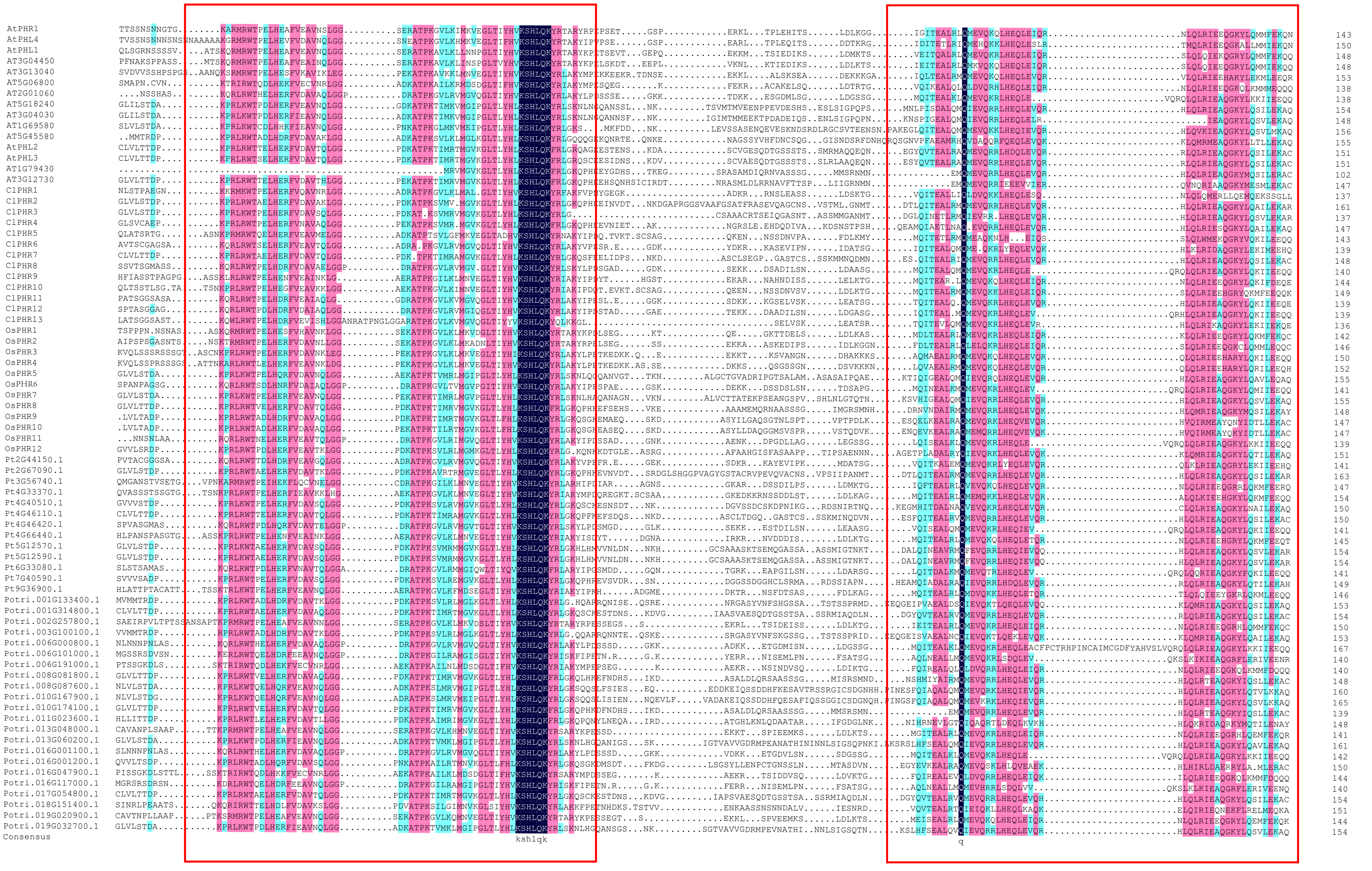


**Supplementary Fig. S1. Multiple sequence alignment of the two conserved ClPHRs domains.**

The Myb_SHAQKYF and Myb_CC_LHEQLE domains were marked in the red box. The highlighted color represents the degree of amino acid identity: black represents a sequence identity of 100%; pink represents a sequence identity between 80% and 99%; blue represents a sequence

Supplementary Fig. S2

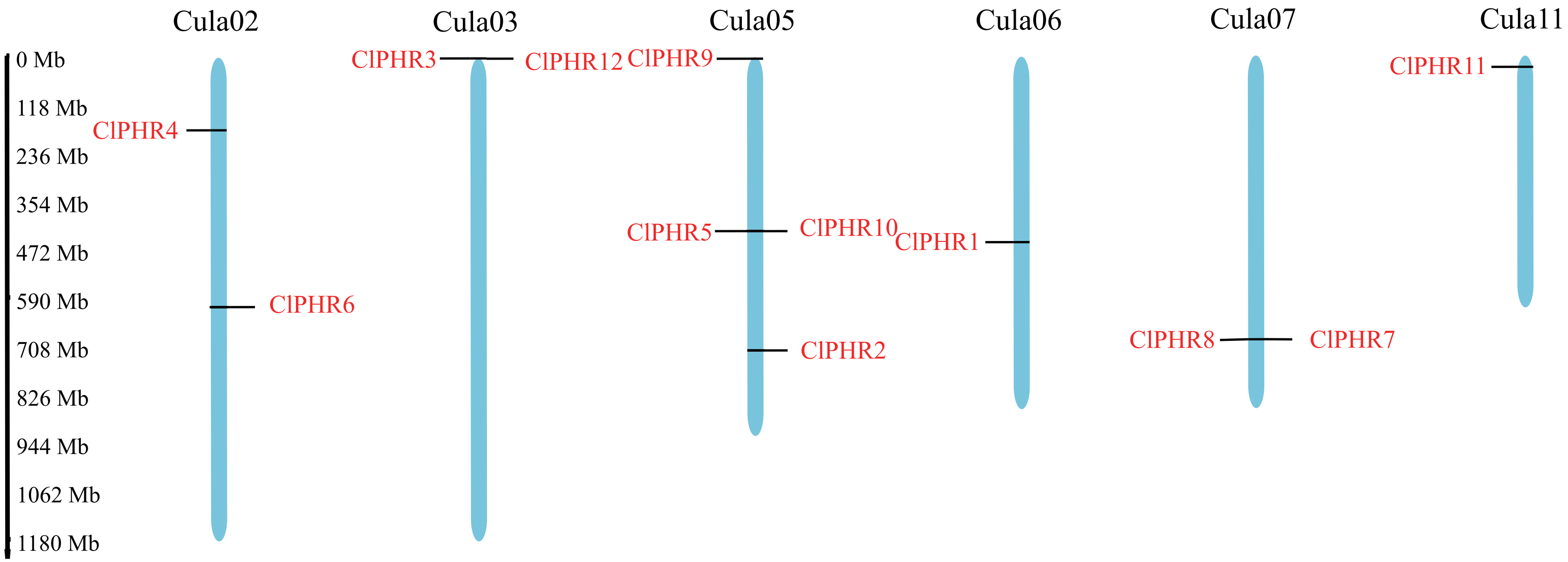


**Supplementary Fig. S2. Chromosomal localization of the Chinese fir PHR genes.**
The chromosome numbers are shown at the top of each chromosome. The name for ClPHR members marked in red color is shown on both sides of each chromosome, and its position on the corresponding chromosome is shown on the left.

### Supplementary Fig. S3

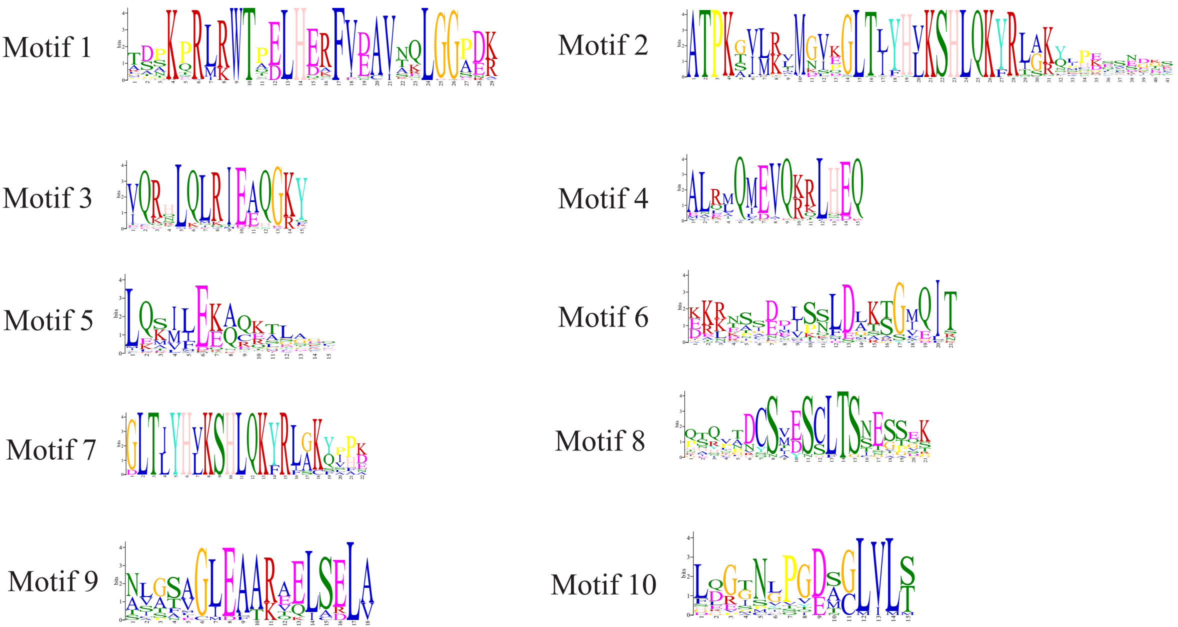


**Supplementary Fig. S3. Schematic representation the sequence logo of the ClPHR protein motif.**

Supplementary Fig. S4

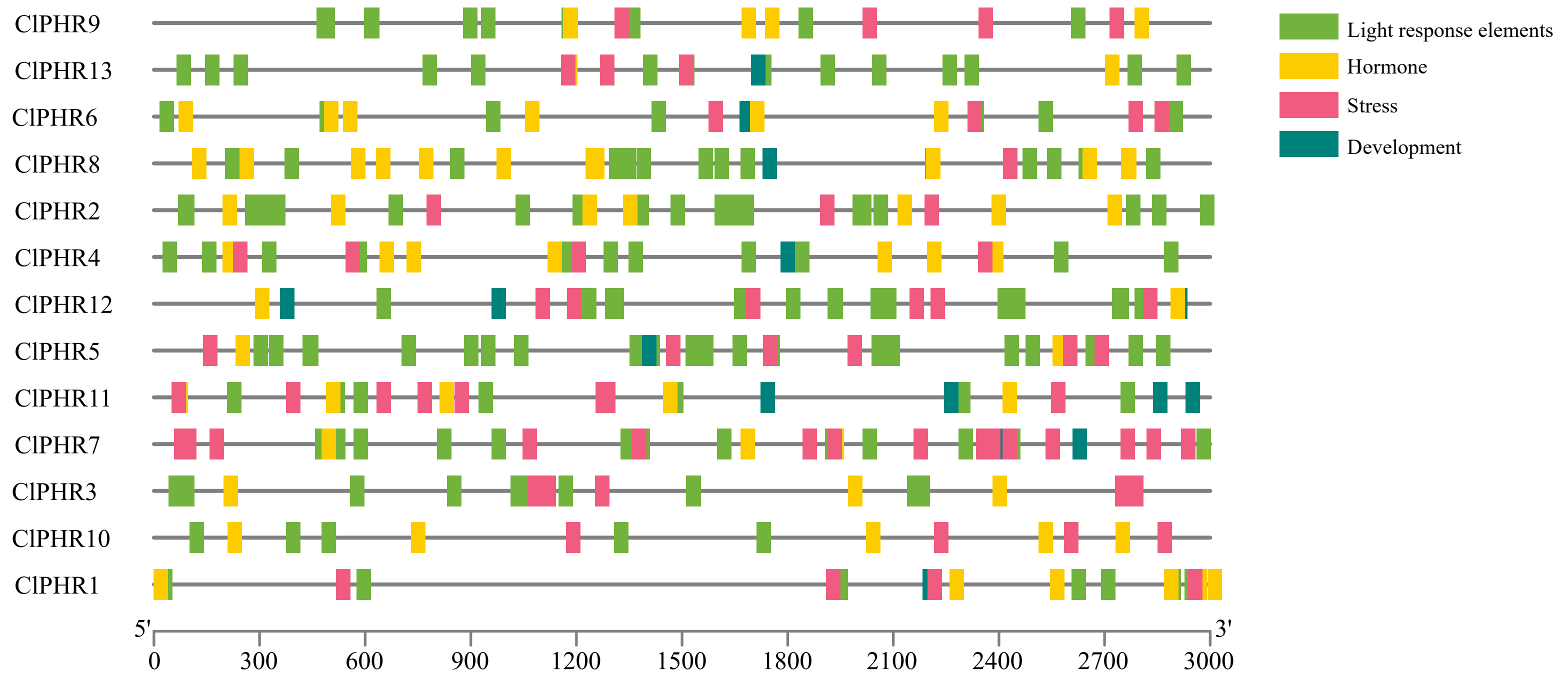


**Supplementary Fig. S4. The Cis-elements of Chinese fir PHR transcription factor.**
The four categories of cis-elements are light-response elements, hormone-responsive, stress-responsive, and plant development. Different colors indicate different cis-elements and their ratios present in ClPHR genes.

### Supplementary Fig. S5

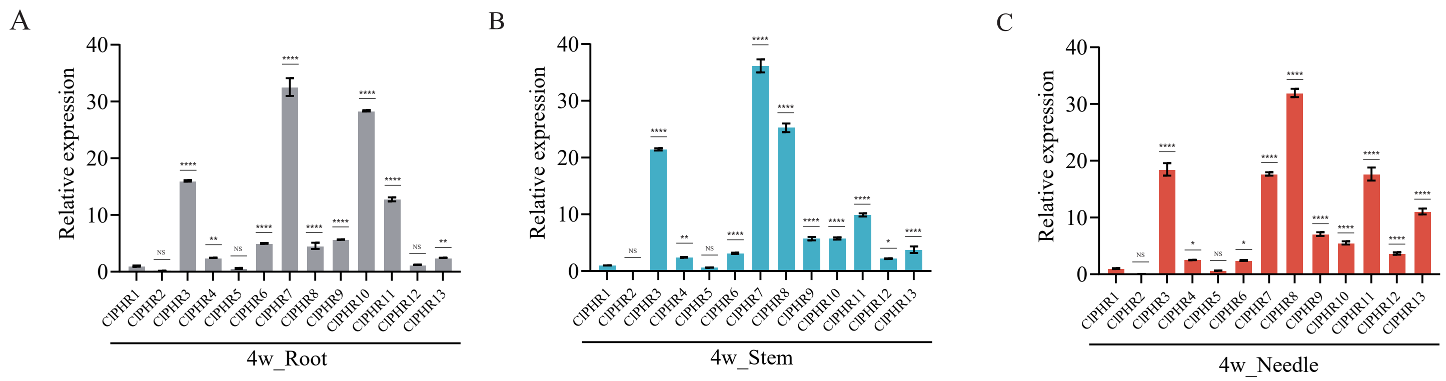


**Supplementary Fig. S5. The relative expression of the Chinese fir PHR gene in roots, stems, and needle leaves.** The relative expression of ClPHRs in the root (A), stem (B), and need leaves (C) of Chinese fir four-week-old seedlings. " ns ", no significant, represents $P > 0.05$," * " represents $P \leq 0.05$ , " ** " represents $P \leq 0.01$ , " *** " represents $P \leq 0.001$ , " **** " represents $P \leq 0.0001$.

Supplementary Fig. S6

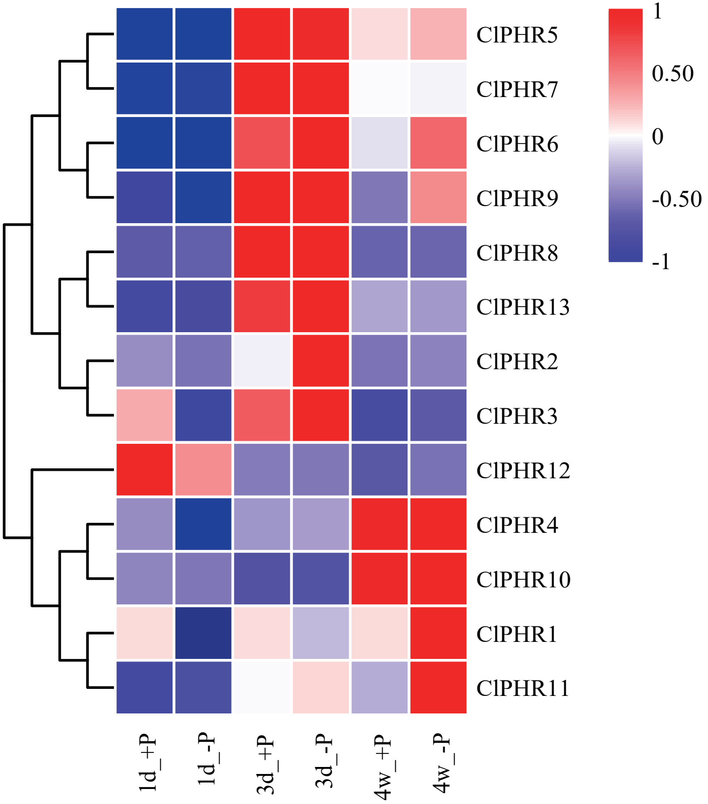


**Supplementary Fig. S6. Schematic heatmap represents relative expression of Chinese fir PHR genes upon Pi deficiency.**

The heatmap represents the expression of Chinese fir PHR genes under 1-day, 3-day, and 4-week Pi deficiency or sufficient treatment.
